# Naïve Bayes Assessment of Mediterranean Basin Ecosystem Wildfires Associated With Vegetation Vigor in August 2021

**DOI:** 10.1101/2022.01.29.478296

**Authors:** Andri Wibowo

## Abstract

The Mediterranean basin, located in the Southern Mediterranean Sea in the North of Africa, is one of the ecosystems that are vulnerable to wildfire hazards. A massive wildfire occurred in this ecosystem in August 2021. Here, this study aims to detect the burnt areas and assess the ecosystem covariates related to the wildfire occurrences. Wildfire assessments were conducted using remote sensing (RS) analysis. Measured ecosystem covariates include vegetation cover, moisture, and the Normalized Difference Vegetation Index (NDVI). The RS analysis confirms at least 17090 ha of vegetation cover were damaged. The burnt vegetation cover was associated with the vegetation moisture and NDVI. The result shows 61.93% of burnt areas were observed within areas that had low vegetation moisture. While 51.75% of burnt areas occurred within areas that had low NDVI values. Naïve Bayes classifier confirms that vegetation moisture and NDVI covariates can be used to estimate the presences of wildfires with values of Kappa, accuracy, precision, recall, and F1 score are 0.7, 0.85, 0.89, 0.8, and 0.84. The area under (AUC) the receiver operating characteristic (ROC) curve values for vegetation moisture and NDVI covariates were 0.945 and 0.399. This value indicates that low vegetation moisture contributes more to the wildfire presences than low NDVI.

## 1. Introduction

### 1.1 Mediterranean wildfires

In the Mediterranean basin (Rowell & Moore 2000), several million hectares of land are burned on a global scale in every year. Based on estimations, 6000,000 to 8000,000 hectares of land we destroyed per year (Carrega 2008) as observed in landscapes along the Mediterranean’s Northern shore. (Hansen et al. 2013). The Mediterranean ecosystems in terms of climate, terrain, composition, and structure; it is extremely sensitive to fire. As a result, Mediterranean regions have been facing forest fires frequently due to continental climate conditions with hot and dry summers.

### 1.2. Remote sensing based wildfire assessment

Remote sensing technology can be used in the risk estimation, detection, and assessment phases of wildfire management. Remote sensing data, which is spectrally sensitive to surface vegetative characteristics and structure, provides rapid, accurate, and reliable information for post-fire damage analysis. Remote sensing satellites can collect multi temporal data and provide synoptic viewing according to Somashekar et al. (2009).

Although various image processing methods are used in burnt area mapping, the main goal of these methods is to determine changes in the reflective characteristics of the vegetation using their spectral signatures during the pre- and post-fire periods. Previous research by Van W agtendonk et al. (2004) has found a significant change in spectral values during this time period, as well as a significant difference between burnt areas and their surroundings (Lobola, et al. 2007).

### 1.3 August’s wildfires

Since 9 August 2021, several forest fires were affecting the North-East of Mediterranean basin. The most affected areas are the regions of Tizi Ouzou and Béjaïa. These locations were known prone to the wildfires and fires have happened since 2010 and continued in 2012. The available research on wildfire in this region is only several studies by Alganci and Sertel (2010), Belgherbi et al. (2018) and very recent by Curt et al. (2020). While wildfire assessments and information based on the recent 2021’s wildfire events is still limited.

Continuous and rapid assessments of burnt areas and their spatial distribution are required for assessing the effects of fire on landscape and ecosystem, particularly in the fragile Mediterranean basin. This evaluation’s findings will be useful to scientists and local governments during the restoration and rehabilitation phases following the wildfires. The determination of burnt area also provides information on land cover change in ecology, which is important data for post-fire restoration of Mediterranean basin ecosystems.

## Methods

### 2.1. Study area

The study area was an ecosystem and landscape located in Bejaia of Mediterranean basin, South of Mediterranean sea, North of Africa..The research was mainly a remote sensing based study that employs the use of remote sensing analysis.

### 2.2 Wildfire event detections

Wildfire hotspot data for in August 2021 were obtained from Terra/Aqua Satellite using the combinations of VIIRS (Visible Infrared Imaging Radiometer Suite) and MODIS (Moderate-resolution Imaging Spectroradiometer) remote sensing sensors (Kumari & Pandey 2019). The VIIRS sensor has resolution of 375 m per pixel and MODIS is 1000 m per pixel. The use of 2 different sensors aims to obtain more fire hotspot data (Tansey et al. 2008).

The VIIRS was used to cover and detect fire hotspots in small areas and MODIS to cover fire hotspots in large areas (Indradjad et al. 2019). The fire hotspot data is then classified as points by using GIS methods with ArcView 3.2 (Gustiandi et al. 2020). The result is a thematic layer in the form of shp files of fire hotspot distributions sharing the same coordinate and projection with Mediterranean basin land-uses layers.

### 2.3 Burnt area classifications

The wildfire and smoke presences obtained from previous step then were classified. This classification process is important to delineate the burnt areas. In this step, burnt area classification adapts the conventional burnt area classification for Sentinel-2 bands, taking advantage of the wider spectrum of visible, red-edge, NIR and SWIR bands. Following is the equation (Filipponi 2018) to determine the burnt areas:

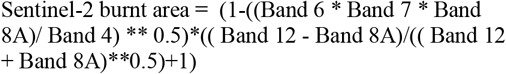

The advantage in the above equation for detecting burnt areas is the use of a band ratio in the red-edge spectral domain, which aim to delineate vegetation properties, combined with a band ratio to detect the radiometric response of the SWIR spectral domain, largely recognized to be efficient in the determination of burnt areas. The burnt areas were used 2 different satellite imageries, one taken on 23^rd^ July 2021 or before the occurrences of wildfires. While another imagery was taken on 27^th^ August 2021 representing post wildfire conditions.

### 2.3 Vegetation cover classifications

The vegetation cover in Mediterranean basin was classified using Geographical Information System (GIS) methods with ArcView 3.2. The method is started with the retrieval of Mediterranean basin boundary and Landsat 8 Operational Land Imager (OLI) images of this basin with a spatial resolution of 30 m per pixel. The Landsat 8 OLI imagery of the basin then classified into several land-uses classes including vegetation cover and barren land covers. The result is a thematic layer in the form of shapefiles (shps) of Mediterranean basin vegetation and barren land covers.

### 2.4 Moisture classifications

Vegetation moisture is displayed using near the infrared (NIR) and the short wave infrared (SWIR) bands in the moisture classification. The SWIR band reflects changes in vegetation water content as well as spongy mesophyll structure in vegetation canopies, whereas the NIR band is affected by leaf internal structure and leaf dry matter content but not by water content. Combining the NIR and SWIR removes variations caused by leaf internal structure and leaf dry matter content, improving the accuracy of determining vegetation water content.

The amount of water present in the internal leaf structure has a large influence on the spectral reflectance in the SWIR interval of the electromagnetic spectrum. As a result, SWIR reflectance is inversely related to leaf water content. In a nutshell, the moisture classification is used to track changes in the water content of leaves and it is computed using NIR and SWIR reflectance’s as follows (Gao 1996):

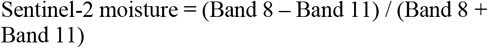

### 2.5 NDVI classifications

In this study, impacted land covers due to the wildfires were assessed using vegetation cover classifications and vegetation indices. The indices were created using spectral values from satellite imagery to perform a quantitative analysis of vegetation health and biomass density. These indices frequently employ photosynthetically active image wavelength portions, such as NIR and Red. The end result of ratio-based indices is an image with pixel values ranging from -1 to +1. The presence of high values in the resulting images indicates that the vegetation is healthy.

The orthorectified pre and post images of the burned area were analyzed using the Normalized Difference Vegetation Index (NDVI) algorithm in this study. The stated algorithm’s formula is as follows:

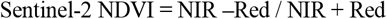

## 3. Results and Discussion

### 3.1. Wildfire event and burnt area detections

Figure 1 depicts the results of wildfire event detections. Based on the figure, there were at least 3 clusters of wildfires producing significant and enormous wildfire smoke plumes. The burnt area classification (Figure 2) has succeeded to distinguish and classify the burnt areas using the wildfires as the references. There are significant differences of conditions between before and after the wildfire events. A massive burnt area was identified after wildfire events have occurred. Since the burnt areas were classified and depicted as a vector object, then it allows to estimates precisely the size of burnt areas. These areas were actually aggregates of several burnt areas that the detail is available in table 1. Based on the calculations, the burnt areas in studied Béjaïa resulted from wildfires have yielded 17090 ha of impacted vegetation covers.

**Table 1.** Burnt area metrics (ha)

| Variable | Sum | Max | Min | St. dev. |
| --- | --- | --- | --- | --- |
| Value | 17090 | 4220 | 0.883 | 424 |

**Fig 1.**
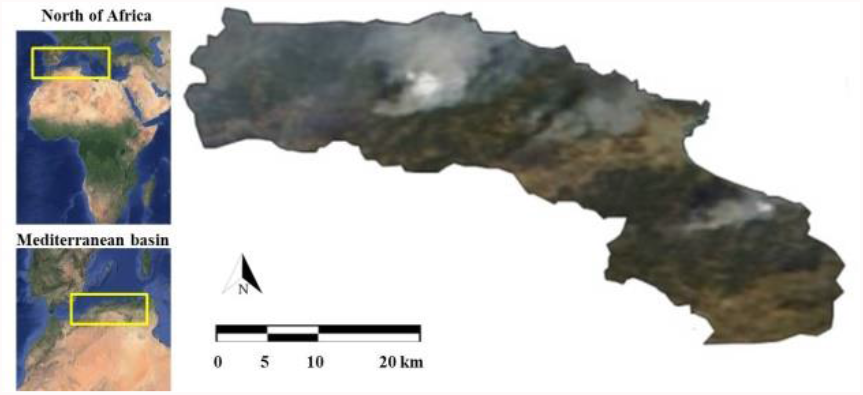
Geographic location of the study areas in Béjaïa landscape of Mediterranean basin, South of Mediterranean sea, North of Africa. Wildfires and smokes started in the study areas as seen by the MODIS Aqua/Terra satellite on 11^th^ August 2021 (insert map: Google Earth).

**Fig 2.**
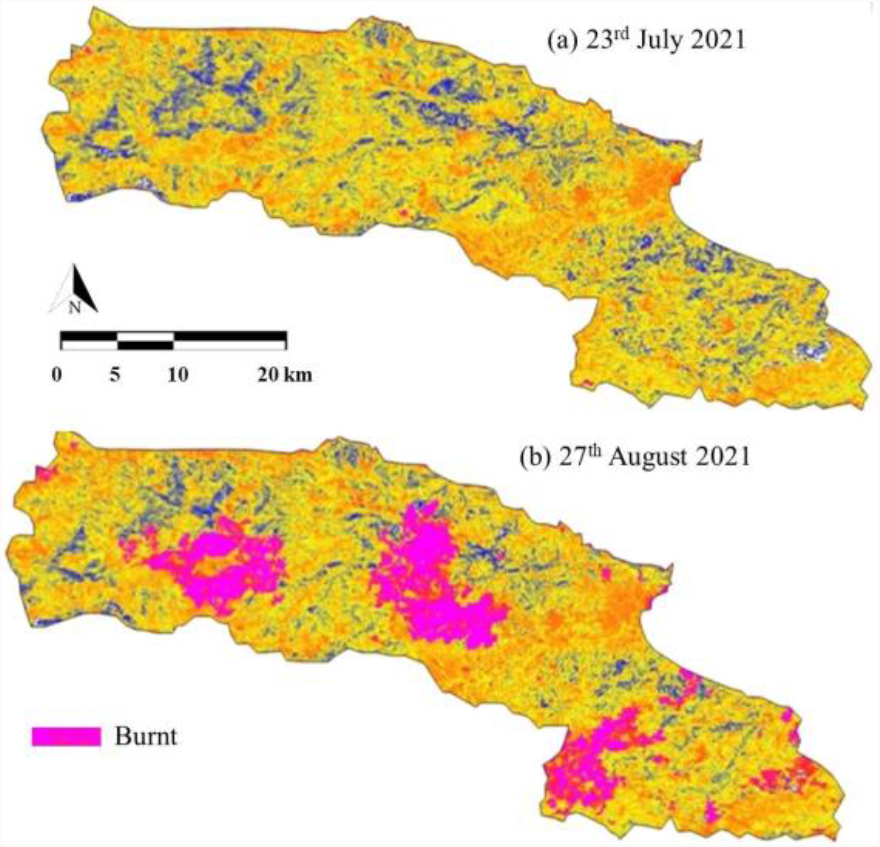
Comparisons of Sentinel 2 imageries taken on (a) 23^rd^ July (pre-fires) and on (b) 27^th^ August (post-fires). Burnt areas were identified using Burnt Area Classification analysis.

In this study, the burnt areas were classified using combinations of wider spectrum of visible, red-edge, NIR and SWIR bands. The current burnt area spectral classification for burnt area mapping specifically designed to take advantage of the spectral characteristics was presented. The present classification benefits from vegetation properties described in the red-edge spectral domains and the radiometric response in the SWIR spectral domain, largely recognized to be efficient in the determination of burned areas. The use spectral information allows to map burnt areas at 20 m resolution and to identify small burnt areas.

### 3.2 Vegetation covariates

The vegetation covariates measured in this study are available in Figure 3. The significant findings are that not all the lands classified as vegetation covered have high vegetation moisture and NDVI values. Some of the vegetation cover was associated with the high moisture and NDVI. In contrast, there are also some of the covers that had low vegetation moisture and NDVI values. While there are land fragments that consistently have low or no cover, low vegetation moisture, and NDVI values. These fragments were usually barren land that had been built permanently to establish settlements.

**Fig 3.**
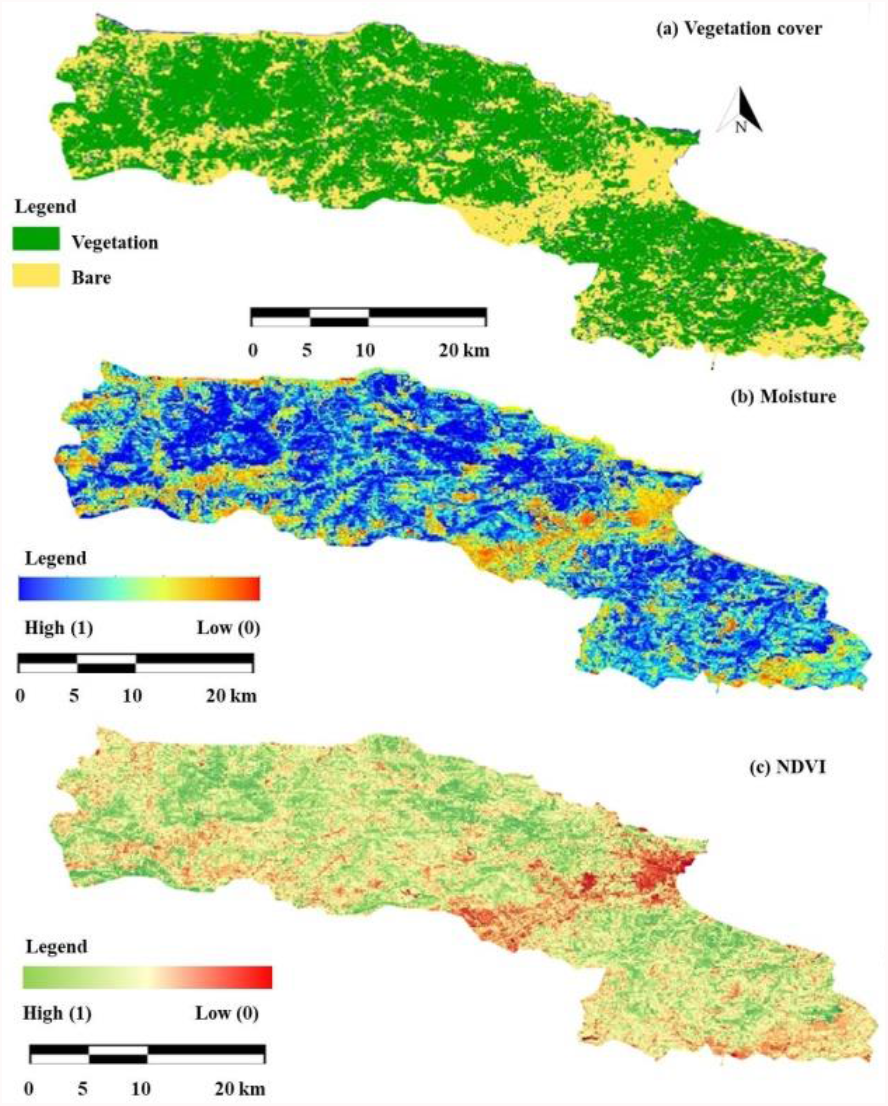
Vegetation cover (a), moisture (b), and NDVI (c) covariates prior to wildfires.

### 3.3 Vegetation covariate and burnt area associations

Classified burnt areas were associated with the vegetation covariates (Figure 4). All burnt areas were overlapped with the vegetation covers. Burnt areas were observed frequently in vegetation covers that have low moisture and followed by the low NDVI values (Figure 5 and 6). Naïve Bayes classifier confirms that vegetation moisture and NDVI covariates can be used to estimate the presences of wildfires with values of Kappa, accuracy, precision, recall, and F1 score are 0.7, 0.85, 0.89, 0.8, and 0.84 (Table 2). The area under (AUC) the receiver operating characteristic (ROC) curve values for vegetation moisture (Figure 7) and NDVI (Figure 8) covariates were 0.945 and 0.399. This value indicates that low vegetation moisture contributes more to the wildfire presences. This finding is in agreement with results from other study. According to Nguyen et al. (2021) in Australia, low soil moisture and high fuel load combined with the occurrence of high temperatures and low humidity in late spring and early summer of 2019 have caused wildfires to start, and the fires have gradually consumed much of the Southeastern coastal areas of Australia where a relatively high fuel load was concentrated.

**Table 2.** Naïve Bayes classifier matrix with overall accuracy of 85% and Kappa value of 0.7.

| Variables | Wildfire |  |
| --- | --- | --- |
|  | Absence | Presence |
| Accuracy | 85% | 85% |
| Precision | 82% | 89% |
| Recall | 90% | 80% |
| F1 score | 86% | 84% |

**Fig 4.**
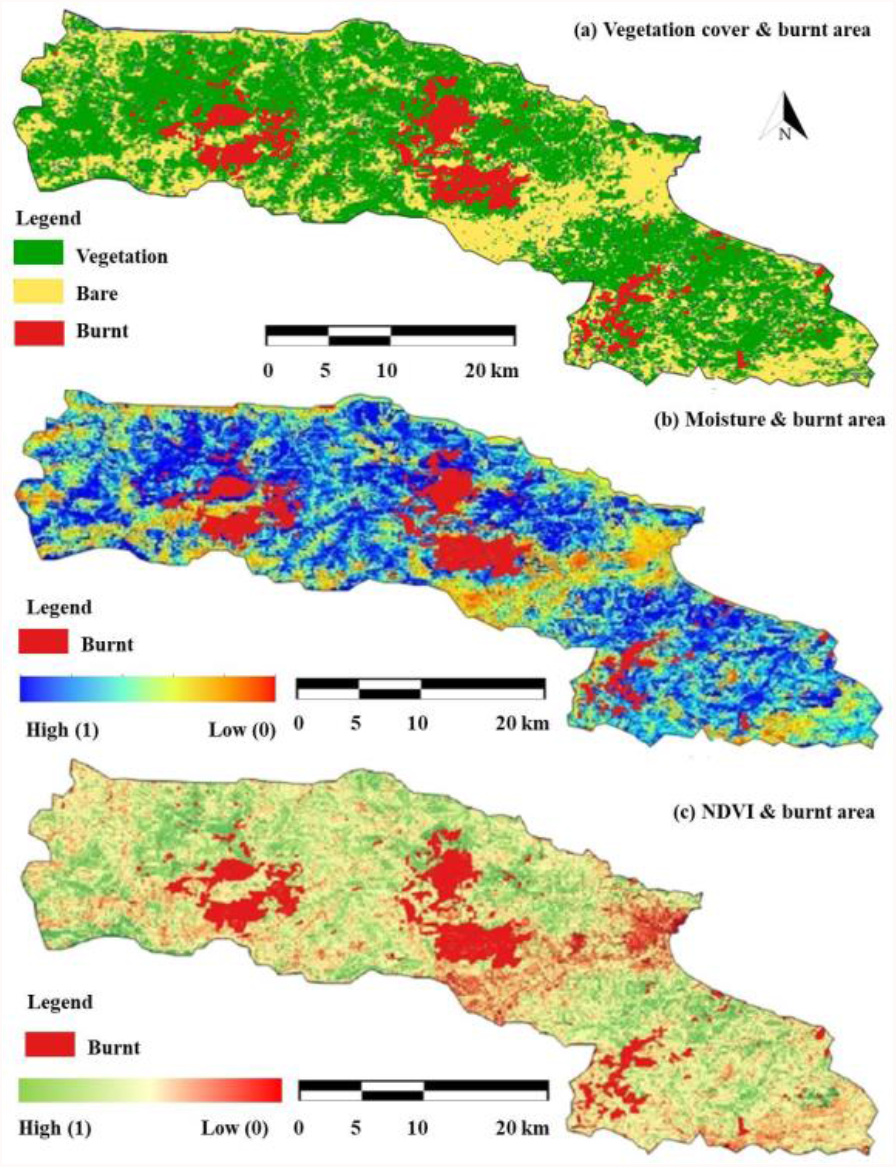
Vegetation cover (a), moisture (b), and NDVI (c) covariates overlaid with burnt areas after wildfires.

**Fig 5.**
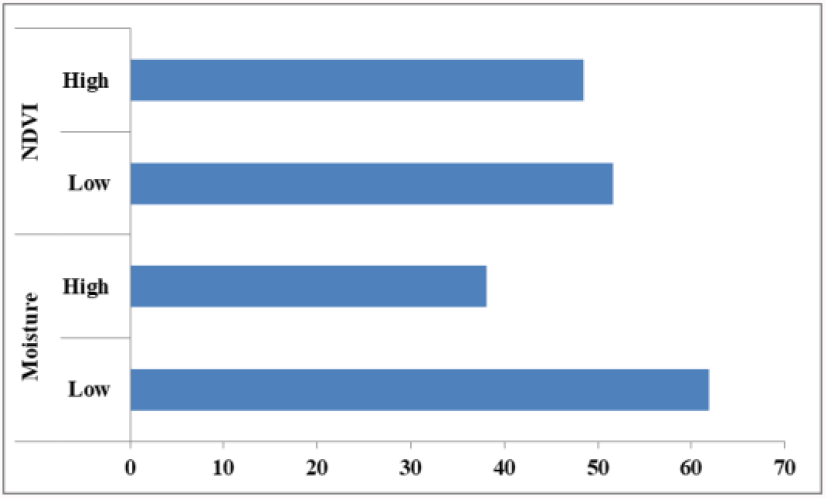
Percentages of burnt areas based on the levels of vegetation moisture and NDVI covariates.

**Fig 6.**
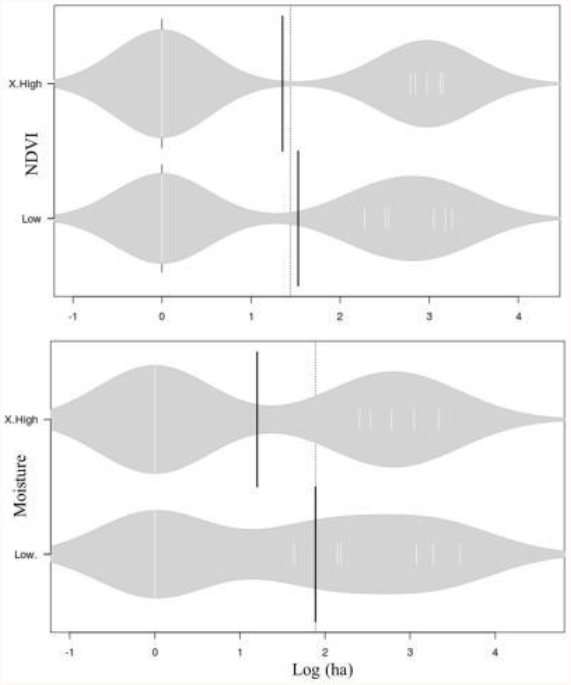
Bean plots of burnt areas (log in ha) based on the levels of vegetation NDVI (top) and moisture (bottom) covariates.

**Fig 7.**
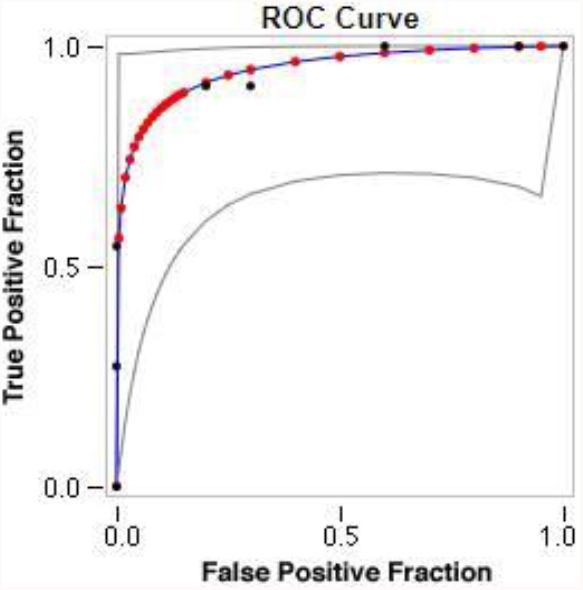
AUCROC for moisture and burnt areas after wildfires with fitted ROC areas of 0.945.

**Fig 8.**
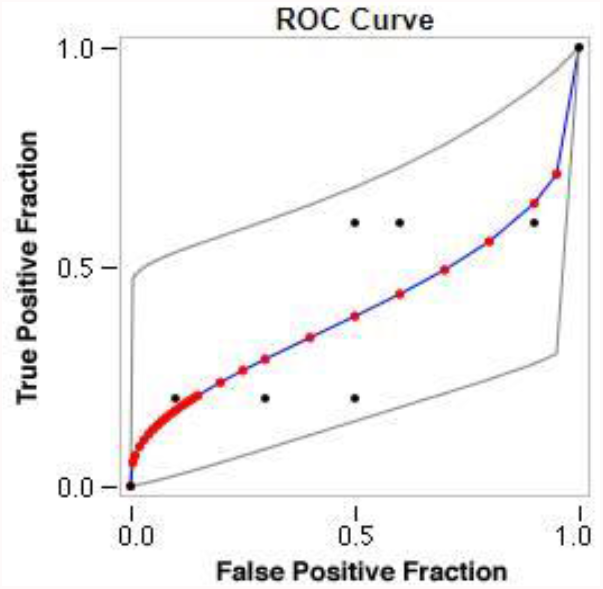
AUCROC for NDVI and burnt areas after wildfires with fitted ROC areas of 0.399.

Occurrences of wildfires within vegetation covers account for 48.42% of burnt areas that have high NDVI in this study indicates high fuel biomass due to the high NDVI that can make the vegetation more prone to the wildfires (Curt et al. 2020). In contrast, occurrences of wildfires within vegetation covers that have low NDVI account for 51.57% were related to the vegetation vigor. Low NDVI indicates declines of vegetation vigor due to the low vegetation moistures (Chéret & Denux 2007). Drop in NDVI value indicates reduced vegetation vigor due to the drop in vegetation moisture content and leads to the vegetation water stress. More vegetation water stress then as the result is an increment of wildfire susceptibility (Sow et al. 2013).

Wildfires occurred in August 2021 were mostly observed in vegetation covers that dominate the hilly areas of the basin landscape. The hilly areas were characterized by slopes. This topography condition is also contributing to the wildfire incidences. From the point of view of biophysics, the expression of fire in a natural space is a function of its environment covariates, including climate, the nature of the terrain, and the fuel present in the area concerned (Pyne et al. 1996). The angle of slope affects the moisture and conservation of the soil, which affects the distribution and composition of vegetation, and thus the characteristics of the fuel and its flammability (Franklin, 1998). In effect, the biophysical factors that influence fire outbreak and spread can have a variety of direct and indirect effects on the fire regime (Whelan 1995).

## Conclusions

To the best of our knowledge, the findings of this study demonstrated that remotely sensed data is an important source for estimating and mapping damaged and affected areas. Furthermore, burnt areas after a wildfire can be easily monitored using multi temporal imagery, which provides important information for understanding the effects of fire on ecology and ecosystem covariates such as vegetation cover, moisture, and NDVI. The GIS environment provides an appropriate platform for understanding and correlating topographic information with satellite imagery, as well as performing analysis for identifying the orientation and distribution of wildfires. The findings of these analyses can be presented as maps and statistical numeric values, which are essential information representation approaches for scientists and policymakers, particularly in Mediterranean basin ecosystems.

